# When Night Becomes Day: Artificial Light at Night Alters Insect Behavior under Semi-Natural Conditions

**DOI:** 10.1101/2023.11.06.565762

**Authors:** Keren Levy, Yoav Wegrzyn, Stan Moaraf, Anat Barnea, Amir Ayali

## Abstract

Light is the most important *Zeitgeber* for temporal synchronization in nature. Artificial light at night (ALAN) disrupts the natural light-dark rhythmicity and thus negatively affects animal behavior. However, to date, ALAN research has been mostly conducted under laboratory conditions in this context. Here, we used the field cricket, *Gryllus bimaculatus*, to investigate the effect of ALAN on insect behavior under semi-natural conditions, i.e., under shaded natural lighting conditions, natural temperature and soundscape. Male crickets were placed individually in outdoor enclosures and exposed to ALAN conditions ranging from <0.01 to 1,500 lx intensity. The crickets’ stridulation behavior was recorded for 14 consecutive days and nights and their daily activity patterns were analysed. ALAN impaired the crickets’ stridulation rhythm, evoking a change in the crickets’ naturally synchronized daily activity period. This was manifested by a light-intensity-dependent increase in the proportion of insects demonstrating an intrinsic circadian rhythm (free-run behavior). This also resulted in a change in the population’s median activity cycle period. These ALAN-induced effects occurred despite the crickets’ exposure to almost natural conditions. Our findings provide further validity to our previous studies on ALAN conducted under lab conditions and establish the deleterious impacts of ALAN on animal behavioral patterns.

**Highlights:**

- ALAN presents a threat for insect populations and biodiversity
- The impact of ALAN on insect behavior is mostly studied under laboratory conditions
- We studied the effects of ALAN on cricket stridulation in semi-natural conditions
- ALAN clearly affected the crickets’ behavior in a light-intensity dependent manner
- The behavioral effects of ALAN were revealed despite the semi-natural environment
- ALAN represents a threat for cricket populations’ fitness

## INTRODUCTION

The daily rhythmicity of light and darkness serves animals for the perception of time (Aschoff, 1954; Borges, 2022) and is the most important environmental cue and *Zeitgeber*, synchronizing internal and external events through entrainment of the animal’s circadian clock mechanism (Aschoff, 1954; Beer & Helfrich-Förster, 2020). Circadian rhythms, as well as daily activity patterns, are manifested in many behavioral patterns, such as locomotion, sleep, foraging, and singing, as well as in physiological processes, including hormonal regulation and gene expression (Beer & Helfrich-Förster, 2020; Rich & Longcore, 2006; Touzot et al., 2022). The behavioral responses to the phases of daylight and darkness depend on the animal’s visual system, its way of life (diurnal, nocturnal, or crepuscular), and the specific nature of the stimulus (Helm et al., 2017; Kronfeld-Schor & Dayan, 2003; Mrosovsky, 1999; E. J. Warrant, 2017). Consequently, the individuals’ synchronization with external environmental rhythms (circadian, circalunar and circannual) is manifested in synchronization of the overall population.

Natural light intensity can vary from 0.001 lx on a clear starlit night, or 0.1-0.3 lx on a full-moon night, to 1,500 lx, and up to 100,000 lx on a cloudy day or in full sunlight, respectively (Rich & Longcore, 2006; E. Warrant & Johnsen, 2013). Artificial light at night (ALAN), the use of anthropogenic outdoor illumination during the night-time, has become prevalent (Falchi et al., 2019; Kyba & Newhouse, 2023) and is increasing worldwide by about 3-6% annually (Hölker et al., 2010). ALAN disrupts the natural light-dark cycle, impacting animals’ daily activity patterns and altering their temporal synchronization (Borges, 2022; Helm et al., 2017; Levy et al., 2021). It has been shown to have severe effects on human health (Zielinska-Dabkowska et al., 2023), on wildlife (Duarte et al., 2023; Rich & Longcore, 2006), and on the environment (Giavi et al., 2020; Jägerbrand & Spoelstra, 2023; Owens et al., 2020; Stewart, 2021). In insects, nocturnal illumination has been reported to induce spatial disorientation (Foster et al., 2021; Owens et al., 2020), resulting in higher predation risk (Davies et al., 2012; Eisenbeis, 2006), increased mortality (Bolliger et al., 2020; Davies et al., 2012; Eisenbeis, 2006; Sanders et al., 2018), reduced pollination (Giavi et al., 2020; Knop et al., 2017), and collectively impeding biodiversity (Falcón et al., 2020).

Laboratory experiments offer a useful tool for ALAN research, enabling controlling and isolating the light stimuli from all other variables. Such experiments, however, lack the complexity of the natural environment including related behavioral aspects such as social interactions. The natural environment incorporates, alongside light-dark cycles, a multitude of rhythmic processes, including thermoperiods, rhythmic soundscape due to the temporal changes in species’ activities, as well as rhythmic intra- and interspecific (e.g., predator-prey) interactions. Ideally, ecological ALAN research would be conducted in the untouched, naturally dark environment, while introducing long-term ALAN manipulations (Hölker et al., 2021). Such field experiments, however, present significant technical challenges. Semi-natural conditions therefore offer an excellent alternative, enabling monitoring the experimental animals under almost natural conditions, while limiting the biotic and abiotic variables by isolating the experimental individuals in dedicated enclosures (Rotics et al., 2011).

Our model organism, the field cricket, *Gryllus bimaculatus,* is a ground-dwelling species exhibiting distinct diel stridulation and locomotion cycles (Fabre et al., 1921; Levy, et al., 2024). Crickets in the laboratory demonstrate temporal shifts in both these activity patterns in response to changes in illumination regimes (Germ & Tomioka, 1998; Levy et al., 2021, 2023; Loher, 1979; Moriyama et al., 2022; Tomioka & Chiba, 1982). Additionally, temporal, circadian variations in gene expression patterns have been observed in the cricket (Levy et al., 2022; Moriyama et al., 2022).

Here we investigated the impact of ALAN on these nocturnal insects in a semi-natural experimental set-up. Individual crickets were subjected to various ALAN intensities in shaded outdoor conditions that incorporated natural temperature and soundscape rhythms. We monitored the individual insects’ stridulation behavior for 14 consecutive days and nights. To the best of our knowledge, this is a first attempt to monitor the behavior of individual insects over such an extended period in semi-natural and thus ecologically-relevant settings. We hypothesized that under such condition, and in contrast to previous laboratory findings, the crickets would maintain entrainment to the environmental conditions and be less affected by the experimental ALAN. Our findings have uncovered a clear deleterious behavioral effect of all examined ALAN intensities in spite of the semi-natural conditions.

## MATERIALS AND METHODS

### Rearing conditions

Crickets were reared in the laboratory under a constant temperature of 26±1°C in a chamber illuminated with white compact fluorescent light (CFL, NeptOn, 6500 K, 380-780 nm, peaks: 547 & 612 nm, Figure S1). The cricket colony was exposed to a 12 h light period of 320 lx (7:00 – 19:00) and a 12 h period of total darkness. Crickets were fed three times a week with dog chow, oats, and vegetables.

### Experimental setup

The experiments took place outdoors, in the grounds of the I. Meier Segals Garden for Zoological Research at Tel Aviv University, from spring until autumn for two consecutive years (2020 & 2021). Male adult crickets (n = 98 results out of a total of 132 individuals) from the indoor breeding colony were housed individually in six separate, transparent enclosures (20x12x13.5 cm each) attached to metal poles at a height of 1.3 m above the ground, 285 cm apart from each other. Each enclosure was lined with a layer of cocopeat, contained an egg carton for shelter, and was sealed with a metal mesh, allowing the cricket to experience natural lighting, temperature rhythms, and soundscape, while preventing its escape or predation. Each enclosure was equipped with a temperature logger and a Swift autonomous sound recording unit (Koch, 2016, Cornell Lab of Ornithology) placed 15 cm above the mesh (Figure 1A&B). Artificial lighting was supplied by CFL light bulbs of the same spectrum described above, but of varying intensities, suspended above each enclosure (Figure 1A&B). Each light bulb was wrapped in aluminum foil to provide the required specific illuminance intensity at each enclosure, and to avoid light emission into the surrounding area (neighbouring enclosures and natural habitat). The enclosures were shaded from direct sunlight by means of black cardboard affixed to the back of the poles (Figure 1A&B), preventing overheating and maintaining similar temperatures in all the enclosures (Figure S2). The different treatments, as detailed below, were conducted simultaneously, maintaining the variation in natural conditions higher between replicas of the same treatment than between different treatments. Crickets were supplied with dog chow and vegetables.

**Figure 1:**
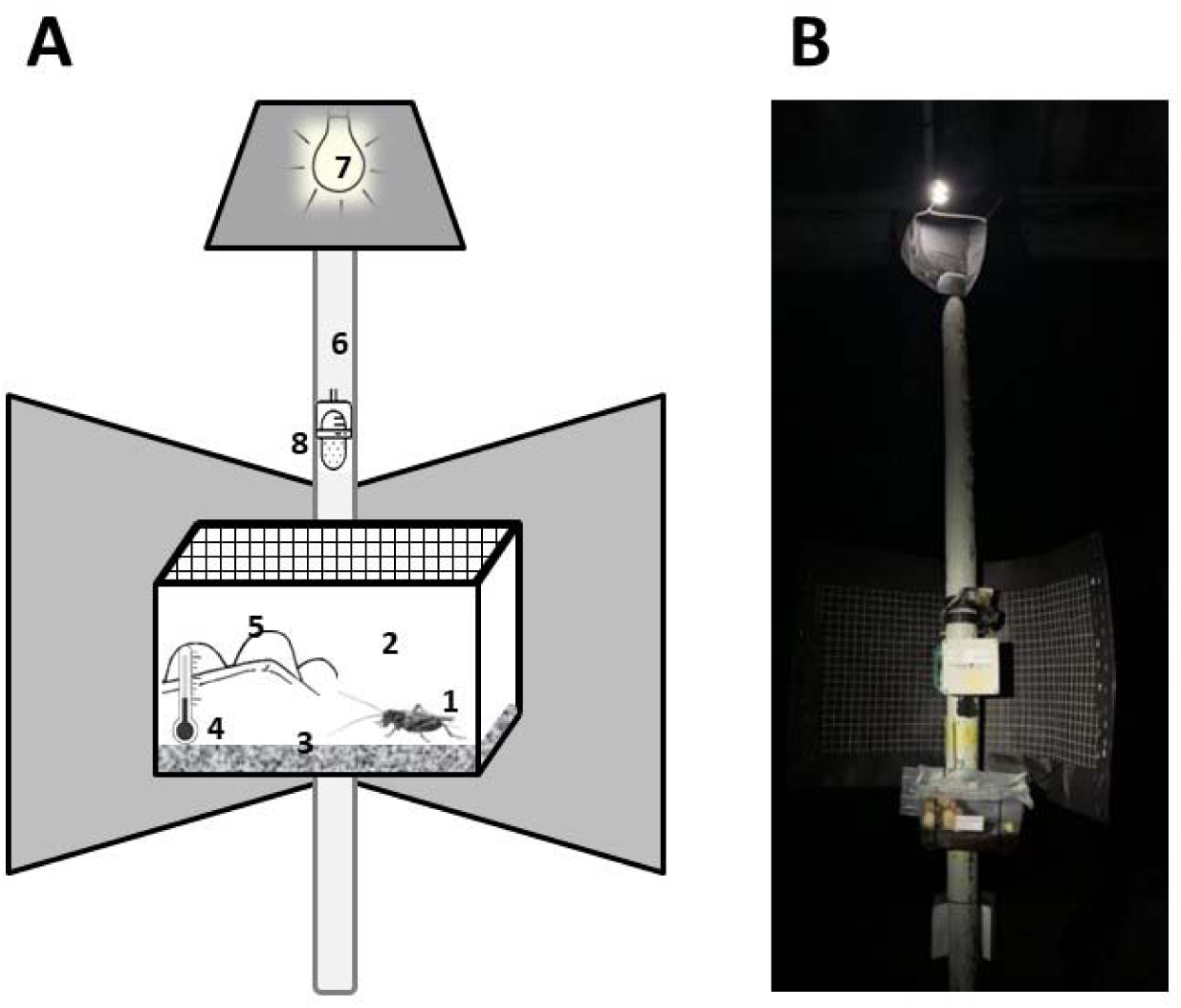
The set-up of the semi-natural ALAN experiment: (**A**) A cricket (1) was placed within a shaded, transparent enclosure (2), lined with cocopeat as substrate (3), a temperature logger (4), and an egg carton (5), attached to a metal pole (6), as well as an ALAN light source (7), and a Swift autonomous sound recording device 15 cm above the enclosure (8), which continuously recorded stridulation behavior. (**B**) The 5-lx ALAN treatment.

### Experimental illumination conditions

All the experimental crickets were exposed diurnally to natural, shaded lighting conditions of a maximum of 1,500 lx, measured at noon across several days using a digital light meter (TES-1337, TES, Taiwan). All light intensities (diurnal and nocturnal) were measured at the four upper corners of the enclosures using the same light meter as above, and then averaged. Each cricket was subjected to one of the following seven treatments, differing in nocturnal illumination: (i) LD (control): dark nights (light < 0.01 lx), (ii) LA_2_: 2 lx, (iii): LA_5_: 5 lx, (iv): LA_15_: 15 lx, (v): LA_100_: 100 lx, (vi): LA_400_: 400 lx, (vii): LL_1,500_: 1,400-1,500 lx (constant light; LL). All light sources were constantly lit. As the light intensity in treatments (i) to (vi), the different levels of ALAN exposure, was much lower than that of natural daylight, it was practically unnoticed by the insects during the day, while treatment (vii): LL_1,500_, represented constant light, as its naturally shaded diurnal intensities were similar to the artificially lit nocturnal intensities.

### Behavioral monitoring

Stridulation behavior (activity) under ALAN exposure was monitored using Swift autonomous sound recording units (Figure 1B), at 32,000 Hz, 16 bit, and 30 dB microphone gain. Each individual cricket was recorded for up to 14 consecutive days and nights.

### Data processing and statistical analysis

Stridulation data extraction and processing followed Levy et al. (2021), using ”R”, version v.3.4.1. (R Core Team, 2020), the “Rraven” open source package (Araya-Salas, 2017), and RavenPro1.5 (Bioacoustics Research Program, 2013). All bioacoustic data, aka stridulation events, were manually validated to avoid false positives/negatives and to eliminate other acoustic sources in the semi-natural conditions, such as bird vocalizations. Only stridulation activity patterns containing at least five consecutive days and nights of behavioral data were used for further analysis. Data processing and statistical analyses were conducted in Python version 3.7 (PyCharm, JetBrains), SPSS version 21 (IBM Corp. Armonk, NY, USA), and Prism 8 (GraphPad Software, San Diego, California USA). The number of detected stridulation events was assessed per animal in 10-minute bouts. For rhythmicity and periodogram analyses, values were normalized for each individual by dividing that individual’s combined values by its own maximum value, resulting in an activity index ranging from 0 (no activity) to 1 (maximal activity). Periodogram analyses of the activity cycle periods were determined using the ImageJ plugin ActogramJ (Schmid et al., 2011), while ClockLab (Actimetrics) was used for onsets and offsets of activity evaluations.

A χ2 test was used to evaluate a possible connection between ALAN intensity and the type of stridulation activity patterns (synchronized or free-run). The medians of the periods were compared using a Kruskal-Wallis test with Dunn’s multiple comparisons post-hoc test. Variance comparison was conducted using a one-tailed Brown-Forsythe test for equality of variance, followed by Welch’s correction. Mean onsets and offsets of activity were calculated by subtracting the time of sunset from that of the onset of activity and subtracting the offsets from sunrise, further converting the results into minutes. A Kruskal-Wallis test was used for means comparisons. Spearman’s rank-order correlation was used to assess a possible connection between ALAN and the frequency of stridulation behavior.

The mean acrophase, presenting the peak of the rhythms evaluated over a minimum of 5 days, was calculated for each animal and period using the CosinorPy package (Moškon, 2020). Mean phases as well as circular statistical analyses were conducted using the Oriana software, v. 4 (Kovach Computing Wales, UK) (Kovach, 2011). The Watson-Williams F-Test was used for comparisons among the treatments’ mean arrow angles and the Mardia–Watson–Wheeler test was used for distribution comparisons among treatments.

## RESULTS

### ALAN affected the type of stridulation activity patterns and their relative proportions

Individual crickets inside their outdoor enclosures were subjected to ALAN conditions at intensities ranging from <0.01 to 1,500 lx. The effects of these different ALAN treatments on stridulation rhythms were manifested in different relative proportions of two types of activity patterns: nocturnal synchronized rhythms with periods of 24 h (Figure 2A1); and free-run rhythms with periods deviating from 24 h (Figure 2A2). The different ALAN exposures elicited a significant light-intensity-dependent increase in the proportion of free-run behavior, from 10% in LD to 92% in LL_1,500_ (Figure 2B, χ^2^(6, N=98)=39.91, *p < 0.0001*), affecting overall 60% of the ALAN-exposed crickets (41 out of a total of 68 crickets).

**Figure 2.**
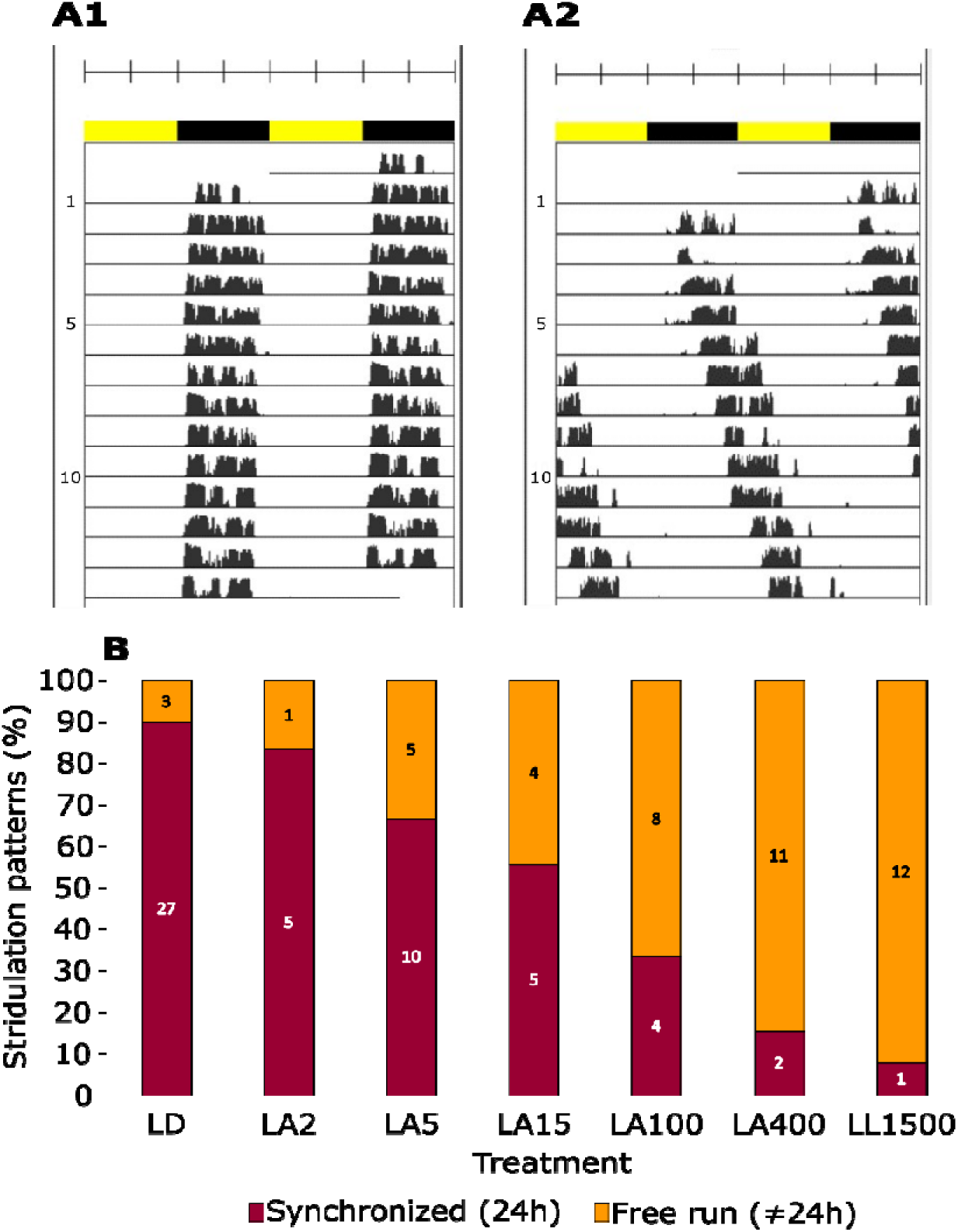
Daily stridulation activity patterns and their relative proportion in adult male crickets exposed to different ALAN intensities under semi-natural conditions: (**A**) Double-plotted actograms representing two rhythmic patterns: (**A1**) synchronized stridulation, with a 24 h period; (**A2**) free-running behavior over a period longer than 24 h (25.7 h in the example shown). Yellow and black bars indicate diurnal and nocturnal phases, respectively. (**B**) Percentage of synchronized (dark red), and free-run (orange) stridulation activity patterns observed in the experimental crickets. The number of synchronized and free-running individual crickets is indicated in the bars (in white and black, respectively; n_tot._=98). Treatments as follows: LD (<0.01 lx, n = 30); LA_2_ (2 lx, n = 6); LA_5_ (5 lx, n = 15); LA_15_ (15 lx, n = 9); LA_100_ (100 lx, n = 12); LA_400_ (400 lx, n = 13); LL_1,500_ (1500 lx, n = 13).

### ALAN altered stridulation activity cycle period in a light-intensity-dependent manner

Analysis of the medians of the period of the recorded behavioral stridulation rhythms revealed light-intensity-dependent differences (Figure 3A, Table S1). While the median period of stridulation activity cycles of the LA_2_, LA_5_, and LA_15_ groups did not differ significantly from those of LD (Figure 3A; Kruskal-Wallis test with Dunn’s multiple comparisons test, p > 0.9 for all; for sample sizes, see Table S1), a significant increase in the period of stridulation activity was evident in all treatments of ALAN ≥ 100 lx (LA_100_, LA_400_, and LL_1,500_; p < 0.02, p < 0.04, p < 0.001, respectively). Furthermore, ALAN also affected the variance of the calculated period: in the ALAN ≥ 100 lx treatments the stridulation activity cycle period displayed a significantly higher variance compared to in the LD (Figure 3A; one-tailed Brown-Forsythe test with Welch’s correction, LA_100_, LA_400_, and LL_1,500_; p < 0.03, p < 0.04, p < 0.01, respectively). The ALAN effects were also manifested in light-intensity-dependent changes in the activity onset and offset relative to the sunset and sunrise, respectively, i.e., the experimental crickets exposed to ALAN ≥ 100 lx delayed both their onset and offset of activity (Figure 3B; Kruskal-Wallis test, for onsets: LA_100_, LA_400_, and LL_1,500_; p < 0.01, p < 0.04, p < 0.003, respectively; for offsets: LA_100_, LA_400_, and LL_1,500_; p < 0.03, p < 0.002, p < 0.02, respectively; see also Table S2). Most conspicuously, the ALAN disruptive effects were accompanied by a discernible trend of an increasing percentage of diurnal stridulation (Figure 3C).

**Figure 3.**
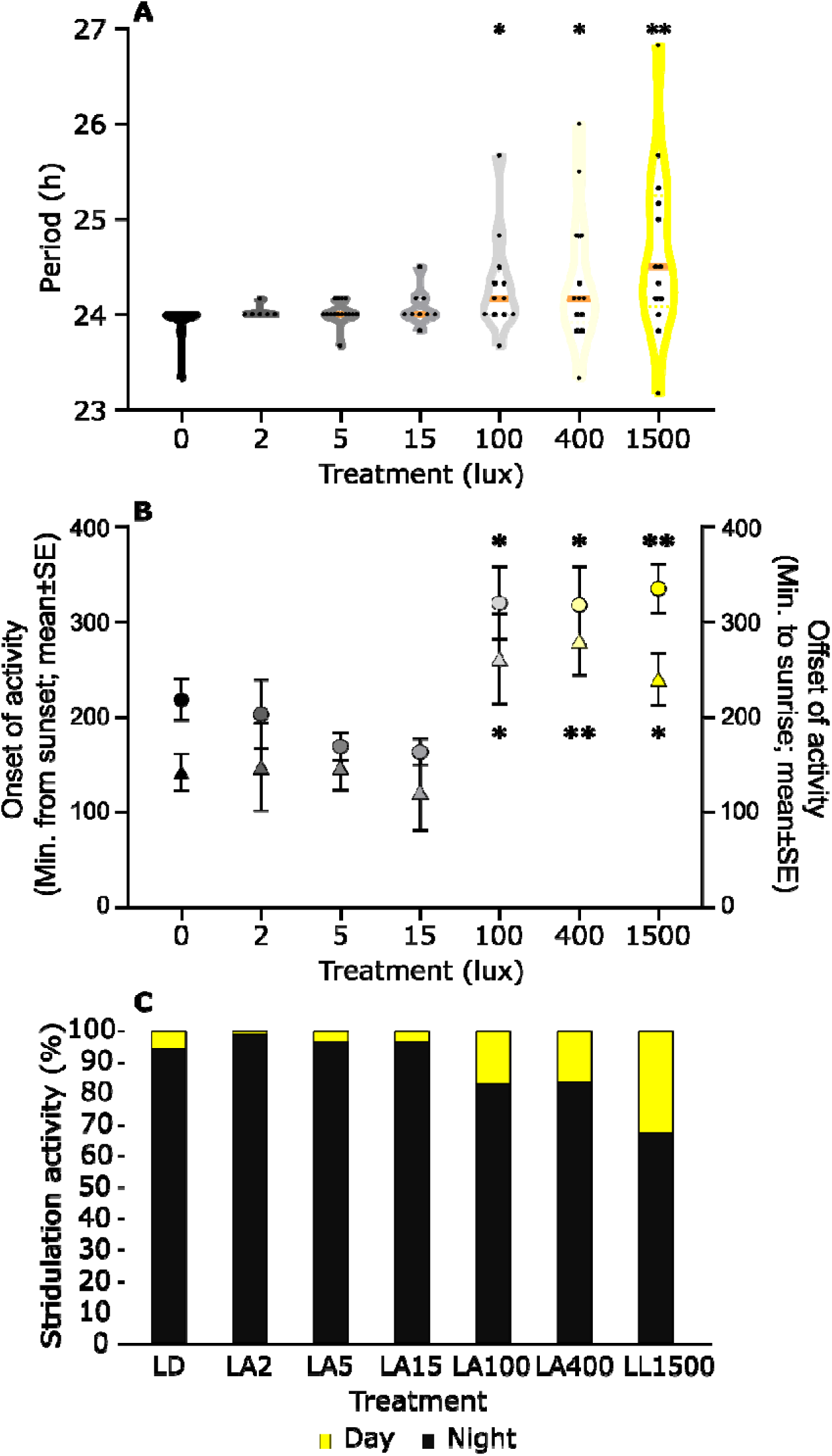
ALAN affects the periods, onset, offset, and timing of stridulation behavior. (**A**) An increase in the medians of the cricket populations’ stridulation periods (medians are colored orange; each individual male is represented by a dot). (**B**) An increase in the mean timespan from sunset to the onset of activity (circles, upper asterisks), as well as from the offset of activity to sunrise (triangles, lower asterisks); and (**C**) in the percentage of diurnal and nocturnal stridulation. n_tot._=98; * *p<*0.05, ** *p<*0.01.

### ALAN did not alter the phase distribution of the stridulation activity

Acrophase refers to the phase of the peak of the fitted Cosinor curve and serves as a convenient phase marker of the behavioral rhythm. Acrophase analysis of the crickets’ stridulation rhythmic cycles, conducted with increasing levels of ALAN intensity (Figure 4, Table S3), revealed no significant changes in the mean timing of acrophase fitted to each individual’s stridulation period (Watson-Williams F-tests, p > 0.1); nor did the phase distribution differ among treatments (Mardia–Watson–Wheeler test, p = 0.37). Pooling all the data together revealed an overall synchronized rhythmicity, expressed in the significant directionality of the mean phase of the population, as well as in a non-uniform distribution (Hotelling’s Test, p = 0.003, and Moore’s Rayleigh Test, p < 0.001, respectively). This indicates an overall resilience of the acrophases of stridulation activity under semi-natural conditions, despite the exposure to ALAN.

**Figure 4.**
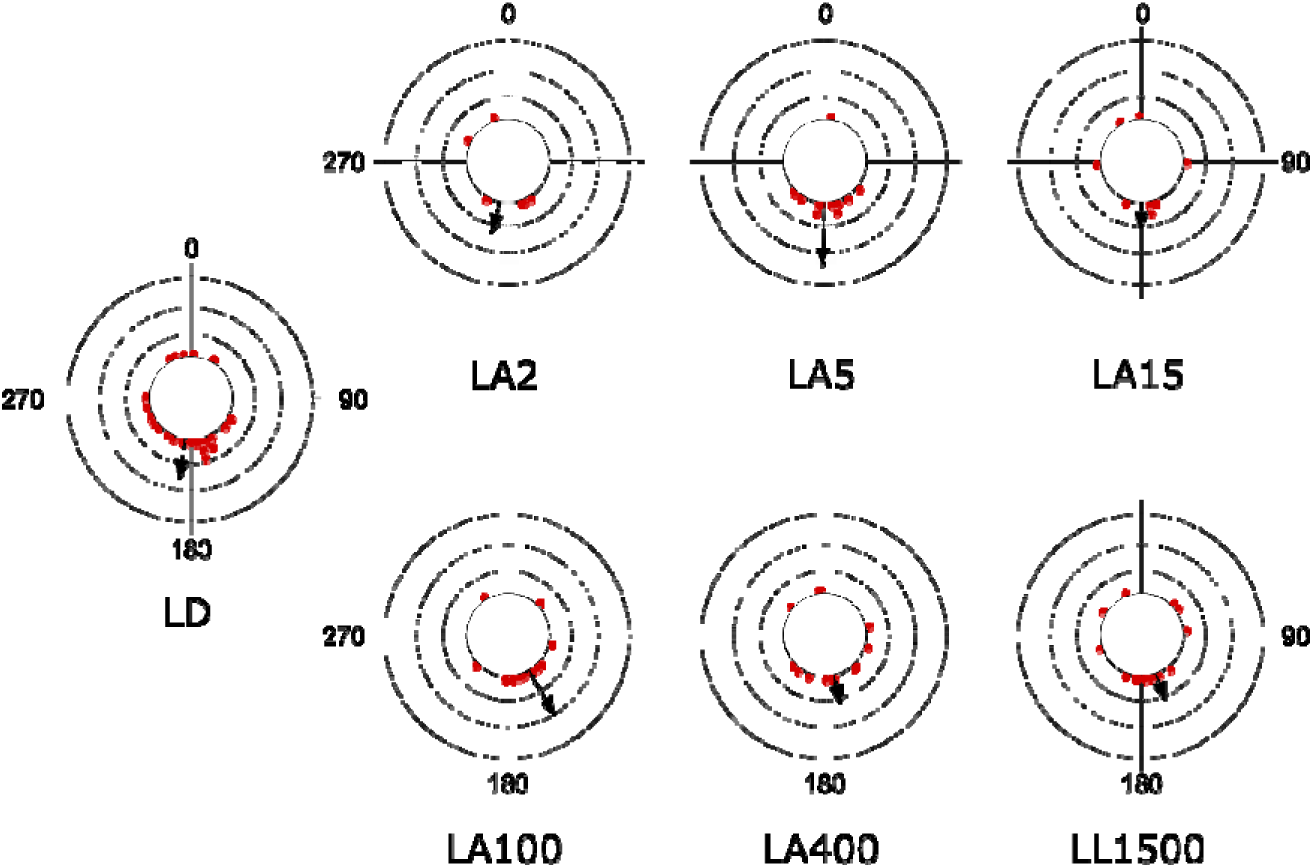
The stridulation acrophase calculations: Under semi-natural conditions, the individual male crickets’ acrophase of stridulation behavior was not significantly affected by ALAN. Each red point in the circular plots represents the mean phase of a minimum of 5 days of stridulation behavior (n_LD_ =30; n_LA2_ =6; n_LA5_ =15; n_LA15_ =9; n_LA100_ =12; n_LA400_ =13; n_LL1500_ =13). Circle grid lines = 1.25. Each black arrow represents the mean vector of the phase, calculated for each experimental group.

### Stridulation frequency is impacted by temperature but not ALAN

The search for possible differences in the stridulation frequencies among the various ALAN treatments yielded no discernible effect (Spearman’s rank-order correlation, *r*_113_ = -0.11, p = 0.26). In contrast, during the experiments which took place over the course of two consecutive years, all the experimental crickets and treatments were subjected to daily and seasonal temperature cycles (Figure S2), which in total were found to significantly affect the stridulation frequency (*r*_111_ = 0.204, p < 0.001).

## DISCUSSION

Crickets serve as prominent model insects for investigating diverse biological and life-history traits. Although studies on communication behavior or population genetics in crickets have occasionally been conducted in the field (Rodríguez-Muñoz et al., 2010; Tregenza et al., 2022), chronobiology research has predominantly been limited to the controlled laboratory environment (Beer & Helfrich-Förster, 2020; Levy, et al., 2024; Tomioka & Matsumoto, 2019). Similarly, investigations into the impact of ALAN on crickets have been primarily carried out under laboratory conditions, with the findings raising significant concerns regarding the impacts of ALAN on the crickets’ overall wellbeing (Durrant et al., 2015; Levy et al., 2021, 2022).

The findings of the current study, conducted under semi-natural conditions and incorporating behavioral monitoring over relatively long time periods, are in general agreement with those of our previous laboratory study (Levy et al., 2021): a light-intensity-dependent effect of ALAN was evident, expressed in the proportion of individuals demonstrating synchronized stridulation behavior. This suggests an ALAN-induced disruption in the population’s behavioral synchronization. Notably, in both the laboratory and under semi-natural conditions impairment of the individuals’ stridulation behavior was observed even under relatively dim ALAN conditions of 2 lx. This sensitivity of crickets to light is not surprising, given that the field cricket *G. bimaculatus* is a nocturnal, ground-dwelling species, with a visual system well-adapted to near-dark conditions (Frolov et al., 2014; Zufall et al., 1989).

While both our previous laboratory study and our current study exhibit similarities in their findings, as described above, they also present several key differences. It is important to acknowledge that under the semi-natural conditions, it was only from light intensities of 400 lx and above that over 80% of the experimental crickets demonstrated a transition towards desynchronized stridulation behavior (Figure 2B). This is in contrast to our earlier laboratory study, where a similar transition occurred already under light intensities of 2 lx and above (Levy et al., 2021). Moreover, another apparent difference between the semi-natural and the laboratory conditions is that, overall, the medians of the activity periods under semi-natural conditions were notably lower (24.0 h and 24.5 h in LD and LL, respectively) compared to those induced by lifelong ALAN exposure in the laboratory (ranging from 24.0 h in LD to 25.67 h in LL (Levy et al., 2021)). Interestingly, differences were also observed in the results of the acrophase analysis. While in the laboratory setting a significant impact on the mean phase vector and phase distribution under LL conditions (Levy et al., 2021) was evident, these same parameters remained stable under the semi-natural conditions (see Figure 4, Table S3).

These differences suggest that the behavior under semi-natural conditions is influenced by additional environmental factors, beyond that of the nocturnal light intensities *per se*. Such environmental factors may include the diurnal light intensity to which the experimental crickets were exposed, and thus also the absolute difference between diurnal and nocturnal light intensities. In addition, sunrise and sunset patterns, and the related changes in illumination spectra, may also have an effect. The potential impact of diurnal light intensities on the susceptibility to ALAN has remained unexplored, to date. However, it should be noted that the diurnal light intensities in the above-mentioned laboratory study were 40 lx (Levy et al., 2021), compared to 1,500 lx in the current semi-natural experiments. It is therefore highly plausible that these differences in light intensity could account for the observed differences in the stability of the activity period medians between the outdoor conditions and the laboratory conditions.

Taken together, our findings from both the earlier laboratory experiments (Levy et al., 2021) and the current semi-natural conditions underscore the crucial impact of ALAN on the cricket as a model insect. Our current study, conducted under semi-natural conditions, emphasizes the importance of field studies that enable multi-sensory modalities to be taken into account in order to comprehensively understand and accurately depict the effects of environmental pollutants on animal populations.

As noted, the experiments presented in this study took place outdoors under near-natural conditions, spanning from spring to autumn, over two consecutive years. Consequently, the experimental crickets were exposed to the natural daily temperature cycle (Figure S2), and to seasonal changes in temperature. Despite this temperature variability, it is striking that 90% of the control crickets maintained a consistent daily activity period of 24 h (Figure 2B, 3A). This finding reconfirms the remarkable phenomenon of temperature compensation (Aschoff, 1981; Saunders et al., 2002), which signifies the stability of the insects’ daily activity period across a wide range of temperatures.

Temperature cycles have been reported to act as an exogenous pacemaker and effective *Zeitgeber* (Aschoff, 1981; Beer & Helfrich-Förster, 2020; George & Stanewsky, 2021) in many animals, including insects. In fruit flies, different clock neurons have been identified as responsible for mediating entrainment by light and by temperature (Yadlapalli et al., 2018). Similarly, crickets that had been made arrhythmic by removing their optic lobes, regained their rhythmic locomotor or stridulatory activity upon exposure to 24-hour temperature cycles (Kannan et al., 2019; Rence & Loher, 1975; Saunders et al., 2002). The peak of the crickets’ stridulation and locomotion behaviors occurred following the transition from high to low temperatures, accompanied by a slightly advanced locomotion activity onset (Kannan et al., 2019; Rence & Loher, 1975). Surprisingly, our findings indicate that over 85% of the crickets subjected to ALAN intensities exceeding 400 lx exhibited free-run stridulation behavior (Figure 2B), despite being exposed to natural temperature cycles (Figure S2). This clear ALAN intensity-dependent effect suggests that the daily temperature rhythms were insufficient to act as a *Zeitgeber* and entrain the crickets’ activity patterns. The fact that, in our study, stridulation behavior was solely synchronized by light conditions and not affected by temperature changes reconfirms light as the most dominant environmental *Zeitgeber,* effectively entraining the circadian clock of adult insects (Aschoff, 1981; Beer & Helfrich-Förster, 2020).

Temperature, however, did affect other aspects of our experimental crickets. Despite experiencing constant temperature conditions in their growth chamber (prior to the experiments), the stridulation frequencies of the individual crickets during the experiments were notably impacted by the outdoor temperature, while remaining unaffected by ALAN. This finding is in accord with previous findings (Doherty, 1985; Symes et al., 2017), suggesting the existence of some degree of plasticity in the stridulation behavior in response to changing environmental cues.

Male field crickets, *Gryllus bimaculatus,* were reported to respond aggressively (acoustically or physically) to male calls within a range of 2 m from their burrows, possibly identifying these as intruders (Simmons, 1988). Furthermore, it was suggested that aggregations of calling crickets serve for intraspecific communication for both females and males, with 2 m being the most frequent distance between two calling males (Simmons, 1988, Figure 2 therein). This pattern of males responding with aggressive or territorial calls to the advertisement calls of other males has also been observed in the cricket *Teleogryllus commodus* (although these did not exhibit positive phonotaxis towards conspecific calls (Thompson et al., 2019)). Given that some insects, such as honeybees and fruit flies, were reported to utilize social cues as *Zeitgebers* for entraining behavioral rhythms (Bloch et al., 2013; Fuchikawa et al., 2016; Lone & Sharma, 2011; Siehler et al., 2021), and since stridulation serves for intraspecific communication in crickets, we hypothesized that our experimental crickets might use conspecific stridulation as a form of social *Zeitgeber*. Although the distance between each of the experimental cricket enclosures in our study was 285 cm (exceeding the 2 m distance noted above), the crickets were still able to audibly detect the stridulation of conspecifics (as evident in the sound recordings). Nevertheless, we found no evidence suggesting synchronization of the experimental individuals according to the cycles of conspecific stridulation. Rather, over 60% of the ALAN-exposed individuals demonstrated free-run stridulation behavior.

Furthermore, the natural environment surrounding the experimental enclosures featured a diverse and daily-cycling soundscape. This included the vocalizations of many bird species exhibiting circadian patterns, traffic noise, and other diurnal human activity. As noted, however, a majority of the crickets exposed to ALAN exhibited free-run behavior, essentially ignoring all these rhythmic environmental acoustic cues. The findings from our study thus demonstrate that the behavior of *G. bimaculatus* crickets is neither synchronized by conspecific stridulation nor by the broader acoustic soundscape. Rather, their behavioral patterns are, as previously suggested, primarily synchronized by light stimuli.

The semi-natural experimental system exhibited several unavoidable yet significant deviations from a fully natural setup. The crickets were isolated from interactions with females and prevented from engaging in reproductive activities, while also not being exposed to any risk of predation. Within their enclosures, these male crickets lacked the possibility to expand or change their habitat or microclimate. Importantly, unlike their free-ranging counterparts in natural habitats that can select less illuminated areas, the experimental crickets were unavoidably exposed to ALAN due to the controlled conditions. These semi-natural conditions, however, effectively mirrored the considerably disrupted environment often encountered in the crickets’ native habitats.

Our findings present evidence of free-run patterns of behavior also within the context of nearly natural conditions (in an illuminated habitat), as well as changes in the timing (onset and offset) of stridulation activity. This desynchronization may impair the crickets’ courtship behavior and potentially put them at higher infection (Durrant et al., 2015, 2020; Westwood et al., 2019) or predation (Zuk & Kolluru, 1998) risk. An increased predation risk arises from two primary factors: first, individuals could become more visually conspicuous against an illuminated background; and second, their acoustic signals could reveal their presence to diurnal predators and parasitoids. For example, certain predatory species, including wasps, geckos and birds, have been documented to have adjusted their visually guided foraging patterns in response to ALAN, expanding or shifting from diurnal to nocturnal activities (Baxter-Gilbert et al., 2021; Dominoni et al., 2013; Gomes et al., 2021). The parasitoid fly, *Ormia ochracea* was reported to be acoustically-orienting to cricket stridulation for locating its host (Zuk et al., 1993; Zuk & Kolluru, 1998). ALAN-induced changes in the timing of stridulation may make crickets more vulnerable to such predators. ALAN was suggested to induce both top-down and bottom-up changes in ecosystems and insect populations (Bennie et al., 2018; Deitsch et al., 2023). Further research is needed to fully elucidate these various and intricate effects.

The present study offers novel insights into the effects of ALAN on insects in semi-natural conditions, and even at low intensities. The growing exposure of large areas to ALAN, commonly observed in industrial zones or due to closely-positioned streetlights, can thus significantly disrupt the insects’ natural behaviors. Our growing understanding of the detrimental effects of the exposure to ALAN, both on the natural behaviors of the individual insect and on the broader dynamics of insect populations, offers a key opportunity for reassessing and reducing the prevalence of outdoor ALAN. Taking proper actions to reduce ALAN would offer a vital step towards the enhanced protection of the natural environment.

## AKNOWLEDGEMENTS

We thank Ronny Efronny for his help with the codes, Mike Pitzrick for his recording advice, Yafit Shachter and Thay Karmin for validations, Yael Lenhardt and Oded Berger-Tal for additional Swift sound recording devices, Michal Seizov Raz and the staff of the I. Meier Segals Garden for Zoological Research at Tel Aviv University for their equipment and assistance with the setup.

## FUNDING

K.L. is grateful to the Cornell Bioacoustics Research Program for supporting attendance at the 2018 Sierra Nevada Sound Recording and Analysis Workshop. We thank the Theodore J. Cohn Research Fund of the Orthopterists Society and The Open University of Israel Research Fund for funding the research.

## AUTHOR CONTRIBUTION

AA, AB, and KL conceived the study and wrote the manuscript; KL and SM carried out the experiments; KL and YW conducted the data analysis.

The authors have no competing interests related to this study.

## DATA AND MATERIAL AVAILABILITY

The supplementary materials and data for this article can be found online at: https://doi.org/10.5061/dryad.6t1g1jx4z (Levy, et al., 2024)

## Supporting information

stridulation of the experimental cricket

---

**Figure S1.**
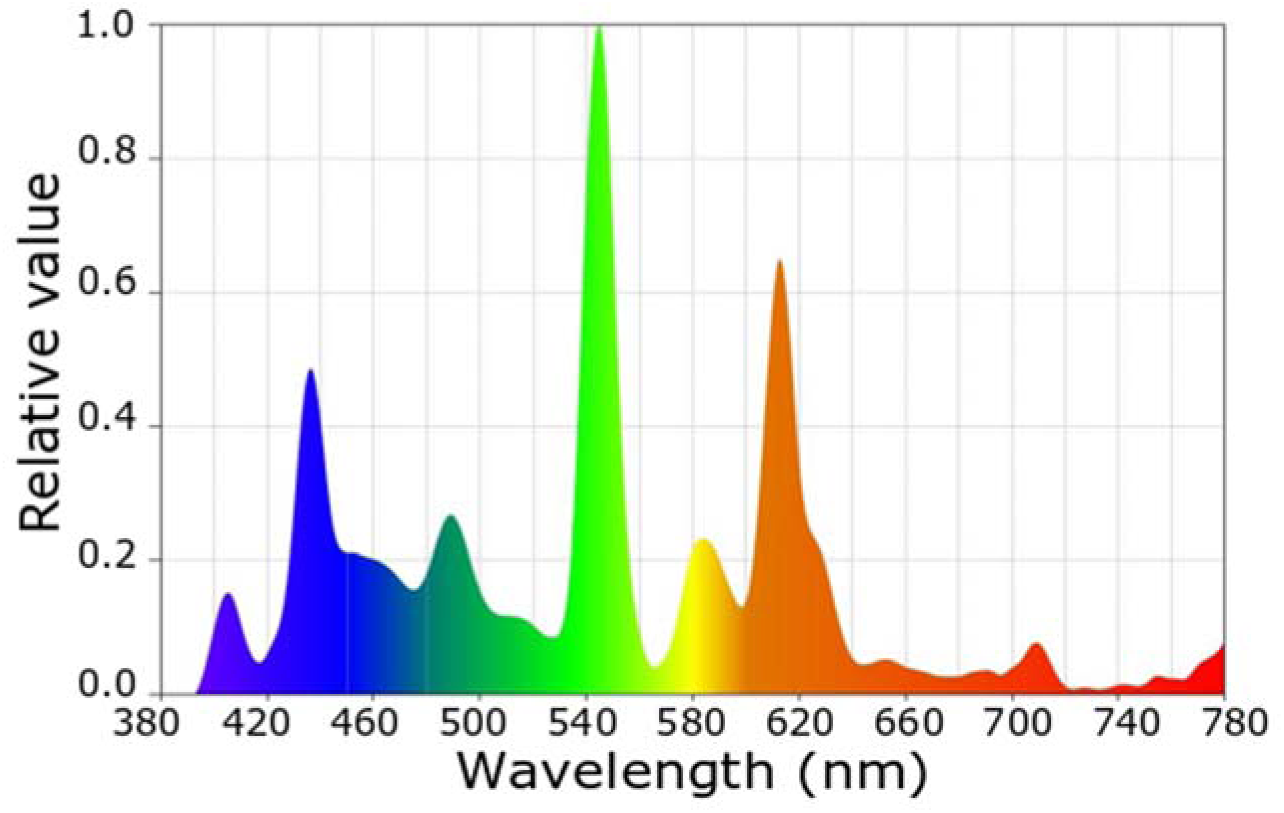
Spectrograms of the white, compact, fluorescent light bulb. (CFL, NeptOn, 6500 K, 380-780 nm, peak: 547 & 612 nm). The light spectra were recorded using a Sekonic Sprectromaster C-700 (North White Plans, NY, USA).

**Figure S2.**
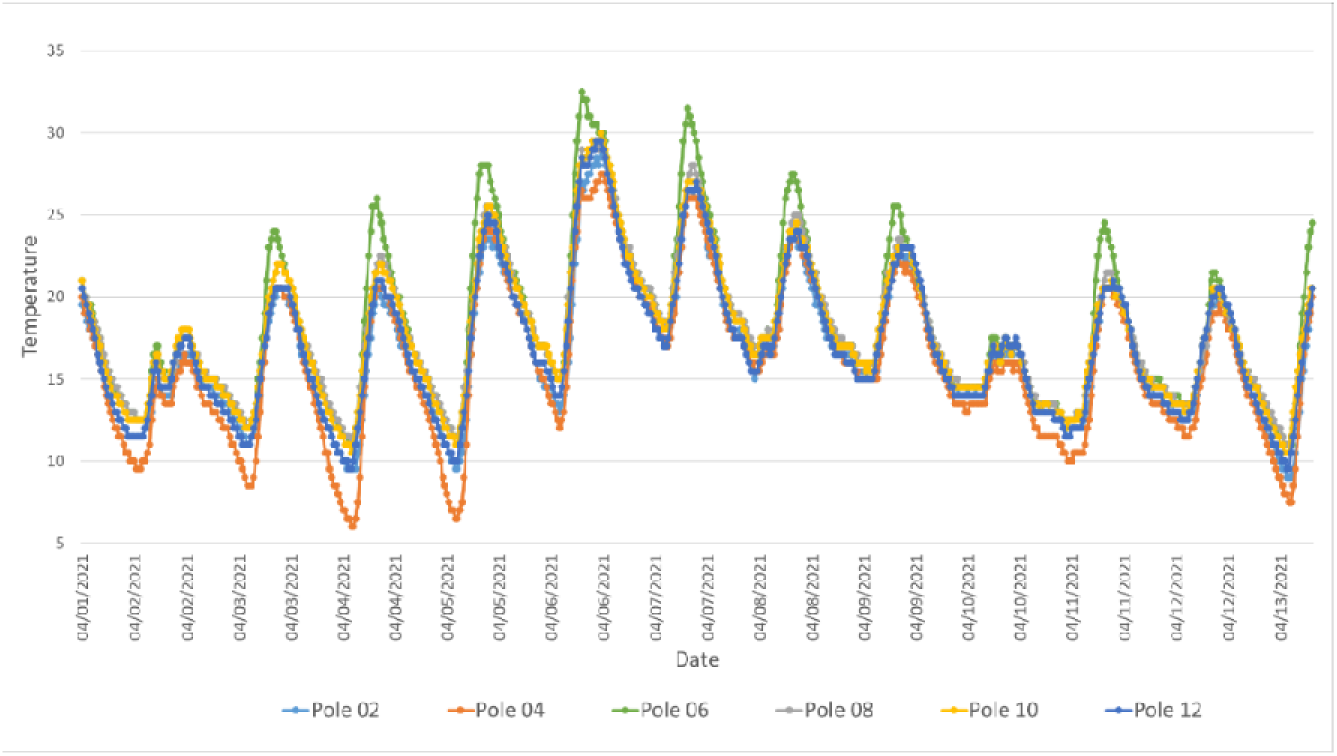
An example of a two-week-long period (01-13/04/2021), representing daily temperature rhythms of the six experimental poles in the semi-natural ALAN experiment.

**Table S1:**
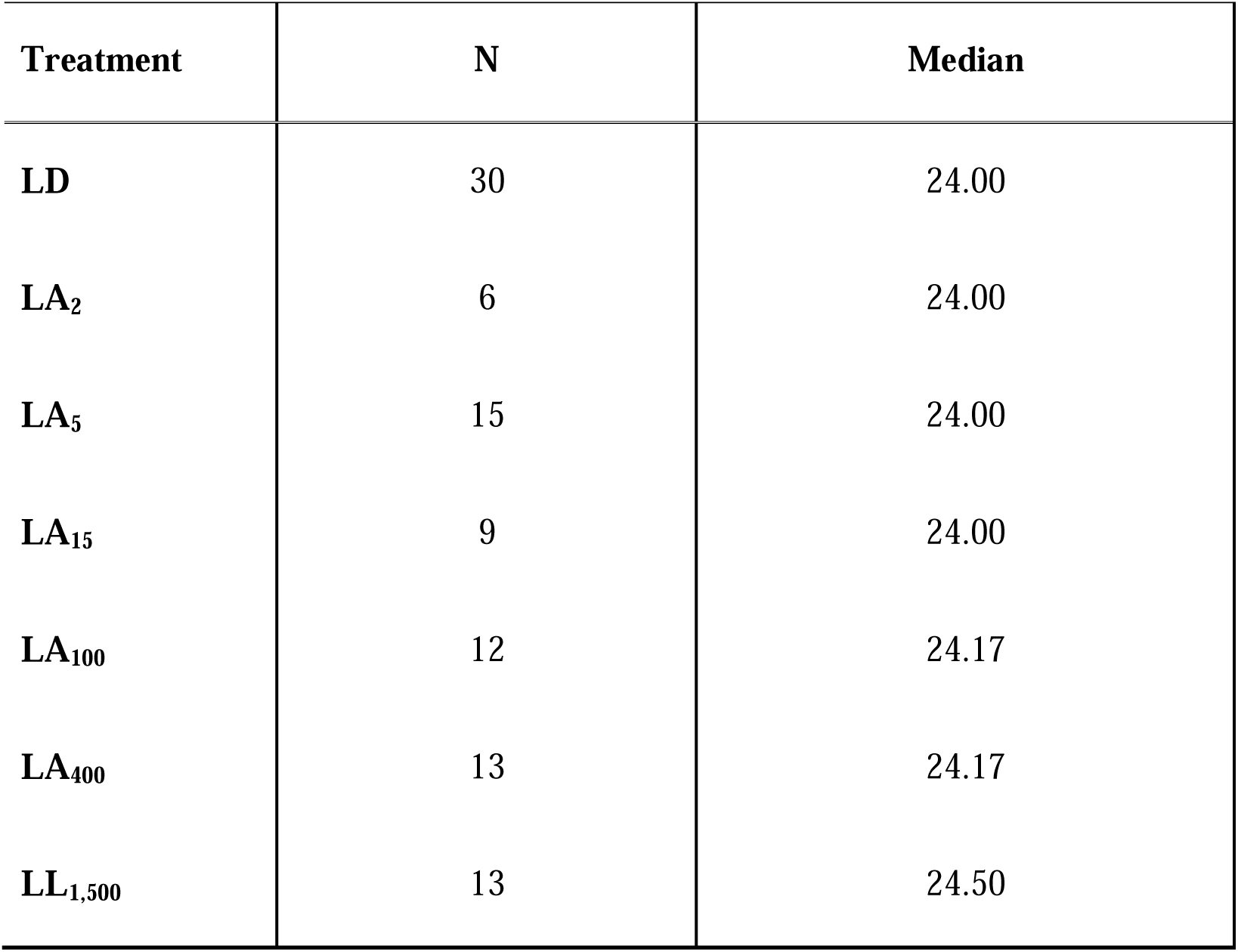
Sample sizes (N) and medians of stridulation activity cycle periods of adult male crickets (*Gryllus bimaculatus*) exposed to seven ALAN treatments under semi-natural conditions. Only individuals with data presenting the activity of more than five days and nights, and with significant activity cycle periods, are included.

**Table S2:**
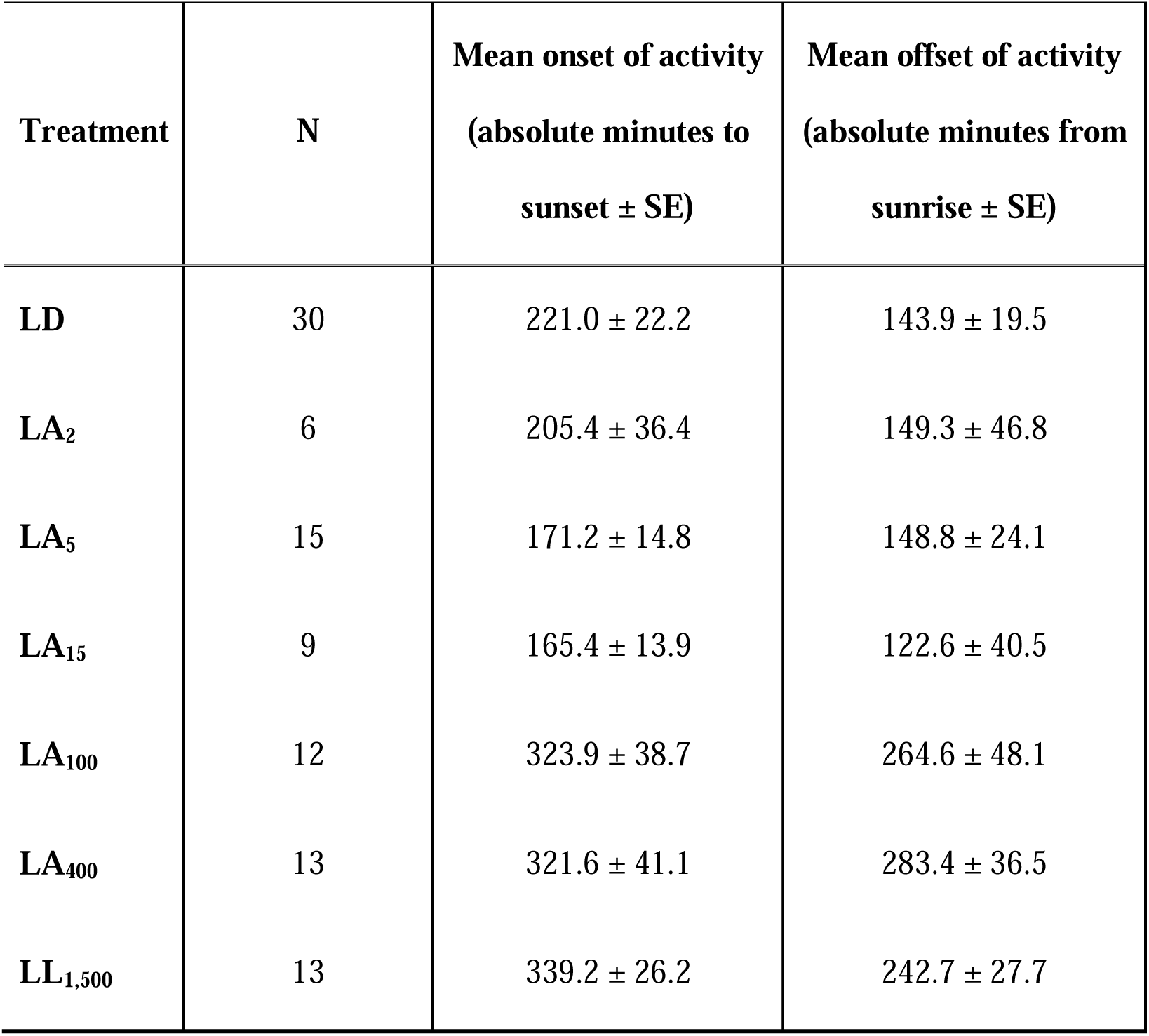
Activity onset and offset analyses of stridulation activity of individual *Gryllus bimaculatus* adult male crickets exposed to seven artificial light at night treatments.

| **Treatment** | **N** | **Mean onset of activity**<br>**(absolute minutes to**<br>**sunset ± SE)** | **Mean offset of activity**<br>**(absolute minutes from**<br>**sunrise ± SE)** |
| --- | --- | --- | --- |
| **LD** | 30 | 221.0 ± 22.2 | 143.9 ± 19.5 |
| **LA<sub>2</sub>** | 6 | 205.4 ± 36.4 | 149.3 ± 46.8 |
| **LA<sub>5</sub>** | 15 | 171.2 ± 14.8 | 148.8 ± 24.1 |
| **LA<sub>15</sub>** | 9 | 165.4 ± 13.9 | 122.6 ± 40.5 |
| **LA<sub>100</sub>** | 12 | 323.9 ± 38.7 | 264.6 ± 48.1 |
| **LA<sub>400</sub>** | 13 | 321.6 ± 41.1 | 283.4 ± 36.5 |
| **LL<sub>1,500</sub>** | 13 | 339.2 ± 26.2 | 242.7 ± 27.7 |

**Table S3:** Acrophase analyses of stridulation activity of individual crickets from the seven artificial light at night treatments under almost natural conditions.

| **Treatm**<br>**ent** | **N** | **Mea**<br>**n (°)** | **Vec**<br>**tor**<br>**Length** | **Medi**<br>**an (°)** | **Varia**<br>**nce (°)** | **Circu**<br>**lar SD**<br>**(°)** | **SE**<br>**(°)** | **95%**<br>**CI**<br>**(+/-)** |
| --- | --- | --- | --- | --- | --- | --- | --- | --- |
| LD | 30 | 188.<br>723 | 0.47<br>3 | 172.4<br>32° | 0.527 | 70.10<br>9 | 14.6<br>93 | 159.<br>919 |
| LA<sub>2</sub> | 6 | 193.<br>394 | 0.37<br>4 | 186.3<br>06° | 0.626 | 80.31<br>9 | 60.8<br>77 | 74.0<br>5 |
| LA<sub>5</sub> | 15 | 181.<br>595 | 0.77<br>5 | 185.4<br>05° | 0.225 | 40.94<br>7 | 11.6<br>21 | 158.<br>814 |
| LA<sub>15</sub> | 9 | 181.<br>461 | 0.33<br>9 | 170.6<br>31° | 0.661 | 84.28<br>8 | 51.1<br>13 | 81.2<br>59 |
| LA<sub>100</sub> | 12 | 149.<br>818 | 0.60<br>8 | 153.3<br>33° | 0.392 | 57.15<br>4 | 17.7<br>12 | 115.<br>095 |
| LA<sub>400</sub> | 13 | 167.<br>94 | 0.34<br>4 | 168.4<br>68° | 0.656 | 83.72<br>2 | 37.5 | 94.4<br>25 |
| LL<sub>1500</sub> | 13 | 160.<br>914 | 0.32<br>9 | 168.4<br>68° | 0.671 | 85.44<br>8 | 40.1<br>78 | 82.1<br>5 |

**Movie S1.**
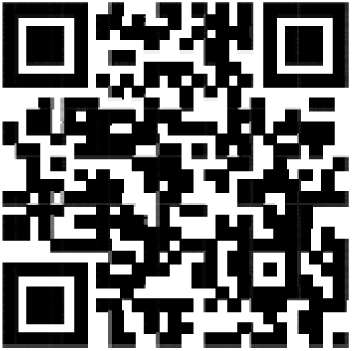
A cricket from the LA_400_ treatment exhibiting diurnal stridulation. Please note the rooster in the background.

## Notes

### Competing Interest Statement

The authors have declared no competing interest.

### Summary of Updates

The methods and discussion sections were improved

