## Supplementary material and tables for "When Night Becomes Day: Artificial Light at Night Alters Insect Behavior under Semi-Natural Conditions"

**SUPPLEMENTARY FIGURES:**

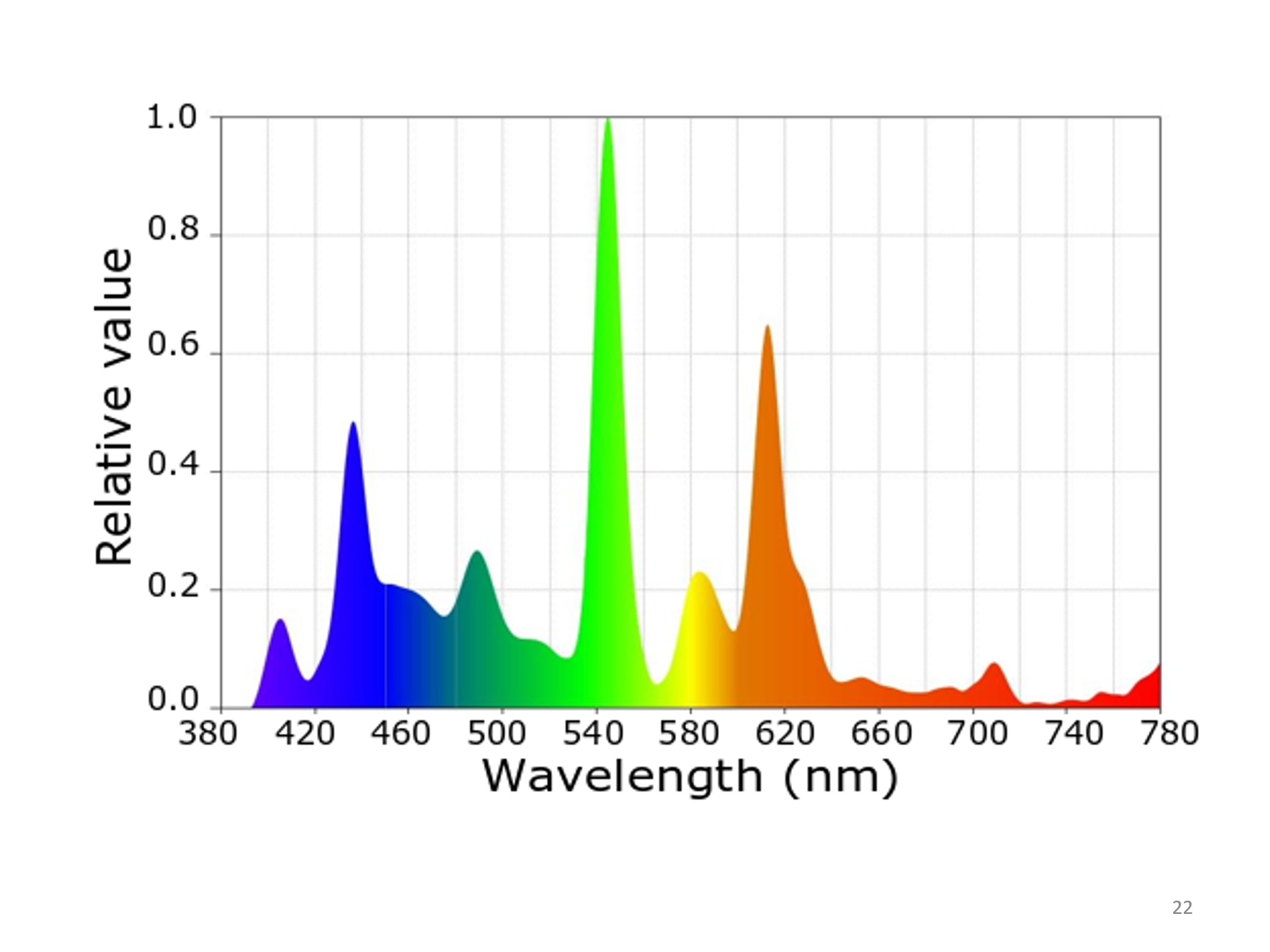

**Figure S1. Spectrograms of the white, compact, fluorescent light bulb** (CFL, NeptOn, 6500 K, 380-780 nm, peak: 547 & 612 nm). The light spectra were recorded using a Sekonic Sprectromaster C-700 (North White Plans, NY, USA).

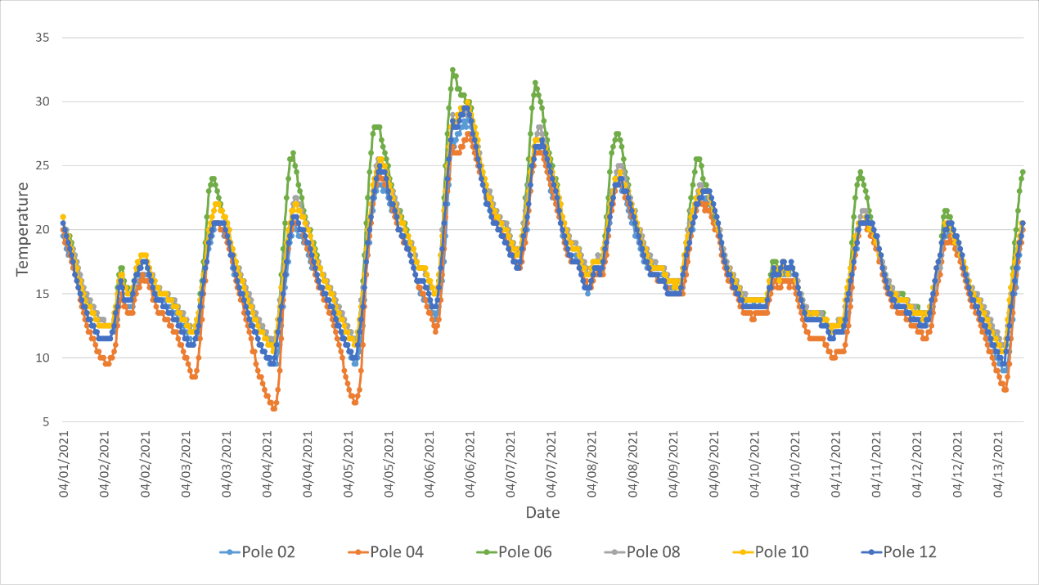

**Figure S2.**  **An example of a two-week-long period (01-13/04/2021), representing daily temperature rhythms** of the six experimental poles in the semi-natural ALAN experiment.

| Table S1: Sample sizes (N) and medians of stridulation activity rhythm periods of adult male crickets (*Gryllus bimaculatus*) exposed to seven ALAN treatments under semi-natural conditions. Only individuals with data presenting the activity of more than five days and nights, and with significant activity rhythm periods, are included. | | |
| --- | --- | --- |
| Treatment | **N** | **Median** |
| LD | 30 | 24.00 |
| LA_2_ | 6 | 24.00 |
| LA_5_ | 15 | 24.00 |
| LA_15_ | 9 | 24.00 |
| LA_100_ | 12 | 24.17 |
| LA_400_ | 13 | 24.17 |
| LL_1,500_ | 13 | 24.50 |

| Table S2: Activity onset and offset analyses of stridulation activity of individual *Gryllus bimaculatus* adult male crickets exposed to seven artificial light at night treatments. | | | |
| --- | --- | --- | --- |
| Treatment | **N** | **Mean onset of activity (absolute minutes to sunset ± SE)** | **Mean offset of activity (absolute minutes from sunrise ± SE)** |
| LD | 30 | 221.0 ± 22.2 | 143.9 ± 19.5 |
| LA_2_ | 6 | 205.4 ± 36.4 | 149.3 ± 46.8 |
| LA_5_ | 15 | 171.2 ± 14.8 | 148.8 ± 24.1 |
| LA_15_ | 9 | 165.4 ± 13.9 | 122.6 ± 40.5 |
| LA_100_ | 12 | 323.9 ± 38.7 | 264.6 ± 48.1 |
| LA_400_ | 13 | 321.6 ± 41.1 | 283.4 ± 36.5 |
| LL_1,500_ | 13 | 339.2 ± 26.2 | 242.7 ± 27.7 |

| **Table S3:** Acrophase analyses of stridulation activity of individual crickets from the seven artificial light at night treatments under almost natural conditions. | | | | | | | | |
| --- | --- | --- | --- | --- | --- | --- | --- | --- |
| **Treatment** | **N** | **Mean (°)** | **Vector Length** | **Median (°)** | **Variance (°)** | **Circular SD (°)** | **SE (°)** | **95% CI  (+/-)** |
| LD | 30 | 188.723 | 0.473 | 172.432° | 0.527 | 70.109 | 14.693 | 159.919 |
| LA_2_ | 6 | 193.394 | 0.374 | 186.306° | 0.626 | 80.319 | 60.877 | 74.05 |
| LA_5_ | 15 | 181.595 | 0.775 | 185.405° | 0.225 | 40.947 | 11.621 | 158.814 |
| LA_15_ | 9 | 181.461 | 0.339 | 170.631° | 0.661 | 84.288 | 51.113 | 81.259 |
| LA_100_ | 12 | 149.818 | 0.608 | 153.333° | 0.392 | 57.154 | 17.712 | 115.095 |
| LA_400_ | 13 | 167.94 | 0.344 | 168.468° | 0.656 | 83.722 | 37.5 | 94.425 |
| LL_1500_ | 13 | 160.914 | 0.329 | 168.468° | 0.671 | 85.448 | 40.178 | 82.15 |

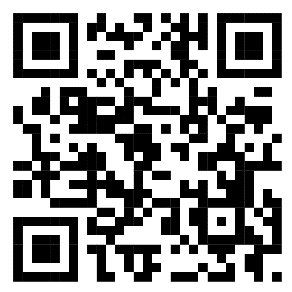
**Movie S1.**  A cricket from the LA_400_ treatment exhibiting diurnal stridulation. Please note the rooster in the background.
